# MLSPred-Bench: ML-Ready Benchmark Leveraging Seizure Detection EEG data for Predictive Models

**DOI:** 10.1101/2024.07.17.604006

**Authors:** Umair Mohammad, Fahad Saeed

## Abstract

Predicting epileptic seizures is a significantly challenging task as compared to detection. While electroencephalography (EEG) data annotated for detection is available from multiple repositories, they cannot readily be used for predictive modeling. In this paper, we designed and developed a strategy that can be used for converting any EEG big data annotated for detection into ML-ready data suitable for prediction. The generalizability of our strategy is demonstrated by executing it on Temple University Seizure (TUSZ) corpus which is annotated for seizure detection. This execution results in 12 ML-ready datasets, collectively called *MLSPred-Bench* benchmark, which constitutes data for training, validating and testing seizure prediction models. Our strategy uses different variations of seizure prediction horizon (SPH) and the seizure occurrence period (SOP) to make more than 150GB of ML-ready data. To illustrate that the generated data can be used for predictive modeling, we executed an ML model on all the benchmarks which resulted in variable performances when compared with the original model and its performance. We expect that our strategy can be used as a general method to transform seizure detection EEG big data into ML-ready datasets useful for seizure prediction. Our code and related materials will be made available at https://github.com/pcdslab/MLSPred-Bench.

## I. Introduction

**E**PILEPSY is a brain disorder that causes repeat unprovoked seizures and afflicts over 3 million people in the USA and over 65 million people worldwide [1], [2]. Seizure symptoms vary from the mild to the extreme including but not limited to shaking, convulsions, and loss of consciousness that may result in serious injury [3]. Electroencephalography (EEG) measures the brain’s electrical activity and can be utilized to mark the start and end of seizures. The prevalence of EEG equipment has resulted in large amount of data that can potentially be used for training machine learning (ML) models for various objectives including detection, and prediction of seizure activity. There are multiple EEG datasets [4]–[7], albeit most of them are either small or have been annotated for detection i.e. labels are attached when seizure is occurring. These benchmarks can be used for training models for detecting seizures. However, an EEG dataset annotated for seizure prediction is currently non-existent.

In general, with enough data wrangling, an EEG data annotated for detection can be transformed into data that is annotated for prediction. A typical seizure comprises four phases: *pre-ictal* (period preceding a seizure), *ictal* (main seiure symptoms occur in this phase), *post-ictal* (recovery period) and the *interictal*. To demonstrate our method, we re-purposed the Temple University Hospital (TUH) seizure corpus (TUSZ) [8] into a corpora that is suitable for predictive modeling. The TUSZ detection data contains 675 patients, 4000 seizures for 10,000 hours, and is diverse for gender, age, channels, and multiple channels. After wrangling the data using our method, we produced 12 different benchmarks with seizure occur(rence) period (SOP) ranging from 1 to 5 minutes, seizure prediction horizon (SPH) ranging from 2 and 30 minutes etc. and resulting in 150 GB of data. This resulting data is suitable for predictive seizure modelling, and is also ML-ready. However, data wrangling requires significant time and effort. In this paper, to make it easier for ML practitioners to develop predictive models using EEG data, we present a methodology and open-source the code necessary to make these benchmarks possible.

### A. Contributions and Organization

This paper has the following contributions in data selection, curation, and annotation to an existing EEG data set:

- Method to transform any EEG annotated data in an ML-ready manner for seizure ***prediction***.
- The resulting method takes data and produces a comprehensive list of benchmarks enabling the training, validation and testing of ***patient-independent*** predictive models.
- Provides an ***open-source*** tool that will enable users to create their own benchmark with any of the EEG data.
- Validates the benchmarks usefulness and quality by ***re-training*** on a previously published deep learning (DL) model.

The paper is organized as follows: Section II briefly discusses the related work, Section III introduces the materials used including the computing resources and the datasets. Section IV discusses the challenges, and Section V describes the methods employed to make the TUH data ML-ready. Section VI shows some of the benchmark datasets created and finally, Section VII discusses the results and concludes the paper.

## II. Related Work

An ML-ready EEG dataset – or a method that can convert data into one - does not currently exist for seizure prediction. One major challenge is that prediction datasets need large amount of gap times between successive (lead) seizures with variable preictal and interictal duration. Most recent state-of-the-art models that apply DL techniques for EEG-based predictive modeling of seizures use the CHB-MIT dataset [9]–[11]. Whereas these works consider real-time implementation, specific models designed for real-time seizure prediction [12], [13] are done on proprietary data from less than 10 subjects. If such models were developed using a far more diverse dataset from a larger cohort, they would be more generalizable [14].

A few works including [5] use the larger EPILEPSIAE dataset which is not available open-source. Other datasets such as Siena [15] and Bonn [6] datasets are either not suitable for predictive modeling or have data from a very small cohort of patients (University of Melbourne [7]). Moreover, the focus as well as annotation of these EEG datasets have been geared towards detection of seizure events. These data sets are clinically useful, but the annotations lack sufficient information about seizures necessary to train ML models for prediction. While TUSZ has a large number of seizures and diversity in the data, the existence of these issues have demonstrated that not having an ML-ready data is a significant bottleneck for developers which explains the absence of detection or prediction models developed using TUSZ data [14] – despite its diversity.

## III. Materials Used

The dataset used to build the seizure prediction dataset is the TUSZ corpus which contains medium to long-term EEG recordings from 675 patients spread across multiple sessions per patient over a period of several years. The python programming language is used to develop the tool including these specific libraries: NumPy for array manipulation and intermediate data storage, MNE library for handling the raw EEG data in European Data Format (EDF) files and the h5py library to store and manipulate the final ML-ready datasets in the “.hdf5” format. All other tasks are accomplished with built-in libraries available as part of the Python 3.8.0 or higher distribution. Let us describe the original TUSZ dataset in more detail.

### A. TUSZ Details

The TUSZ contains data from a total of 675 patients and at the patient-level, the data has already been divided into training, validation and testing. The training set contains a total of 579 patients that recorded 1175 sessions, of which only 208 patients recorded at least one seizure and all seizures are contained within 352 sessions. In contrast, the validation and test sets have a total of 53 and 43 patients of which 45 and 34 recorded at least one seizure, respectively. The seizures were spread over 113 and 63 sessions out of a total of 342 and 128 sessions, respectively, in the validation and the test sets. Overall, data is collected from 675 spread across 1645 sessions of which 287 patients have seizures spread across 528 sessions. This information is illustrated in Fig. 1.

**Fig. 1.**
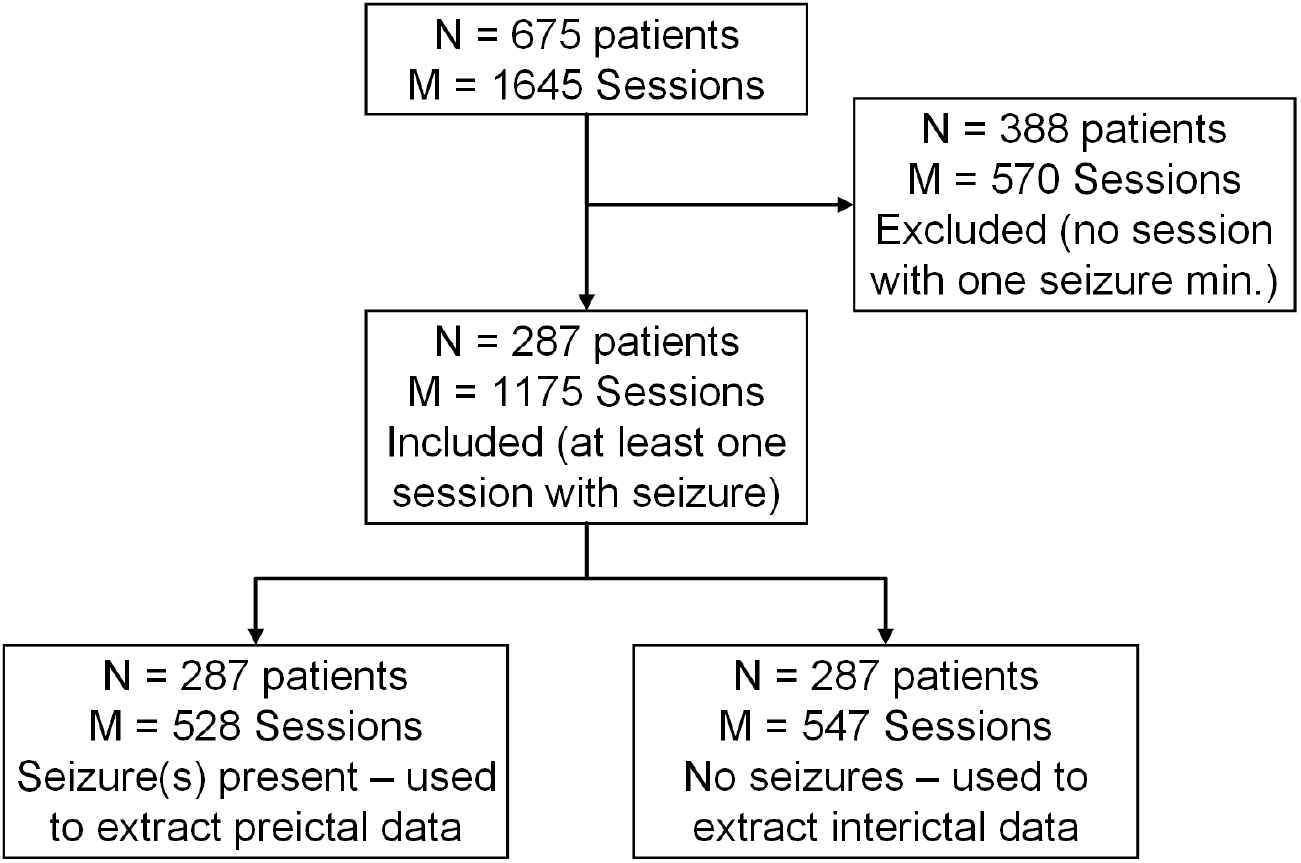
Illustration of how we selected the sessions for extraction of preictal and interictal data.

In total, the 1645 sessions comprise 7377 EEG recordings where multiple recordings belong to the same session due to storage limits on the file size of individual records. The EEG recordings are stored as “*.edf” files. In general, the International 10-20 system was used for electrode placement and data collection. Because the data was collected over multiple years from different equipment by different people, there is a variation in the used sampling rates and the number of channels used. The majority of the sessions used a sampling rate of 256 Hz (5,386) but there were also some sessions that used 250 Hz, 400 Hz, 512 Hz and 1000 Hz. This is taken into account when generating the benchmarks. EEG uses reference electrodes as ‘electrical ground’ to measure the voltage levels. There are two referencing techniques: automatic reference (AR) and linked ear (LE). In the TUSZ, each of those are further divided into two categories as AR1 and AR3, and LE2 and LE4.

## IV. Bottlenecks and Challenges

The goal of this work is to transform multi-channel EEG time-series data into ML-ready data for solving the classification problem of discriminating between preictal and interictal patterns. A successful predictive model will be able to identify preictal patterns or biomarkers with high sensitivity. To accomplish this task, there are several bottlenecks and challenges that need to be overcome related to the data itself and also related to the ML process as detailed in the next subsections.

### A. Different Number of Channels and Reference Types

One bottleneck for ML models is the plethora of different channels and reference types that may be available in a given data cohort. For our purposes, there were four different reference types (AR1, LE2, AR3 and LE4), and for each session, data was available in the range of channels numbering from 25 to 37. Since there are more than 1000 possibilities for the AR type and 50 possibilities for the LE type, this can become a bottleneck due to two reasons. From the data processing perspective, some of the channels are not named with the international 10-20 system format such as P1, O2, T4, etc. Instead, they are labeled with integers where for example, more than 75 channels in various records are labeled with numbers in the range [20, 128]. From the perspective of making data ML-ready, it is imperative to have a uniform number of channels from electrodes located at the same positions of the brain according to the 10-20 system. The reference type is a bottleneck because the same EEG electrode (e.g. P1) may represent a different value for the AR type compared to the LE type, due to having a different electrical ground. Therefore, for our benchmark, we follow the common clinical standard of monitoring epilepsy using montages where each montage is the difference of two electrodes (and their respective references), and eliminates the effects of different types of references. We note that there are 19 useful EEG channels common to AR1 and LE2 while AR3 and LE4 have 17 common channels. To maximize the number of sessions, we use 17 channels common to AR1, LE2, AR3 and LE4 to ensure a uniform montage length of 20.

### B. Useful vs Non-Useful Preictal Durations

Recall that preictal phases are not readily identifiable on an EEG and may vary for different subjects and seizure types [16]. Usually, detection datasets are marked with the start and end of ictal phases. To make the dataset usable for predictive modeling, two new variables need to be defined: the seizure prediction horizon (SPH) and the seizure occur(rence) period (SOP). In our work, the SOP represents the gap time in minutes before a seizure before which prediction should be guaranteed. The SPH is the period during which a seizure must be predicted. In other works, SPH and SOP have been used interchangeably [11], [16]. Let’s assume a seizure starts at time *T*_*start*_, we extract preictal samples from the range [*T*_*start*_ *− SOP − SPH, T*_*start*_ *− SOP*]. Ideally, the preictal period will match the SPH exactly but unlike ictal phases, the preictal phases cannot be marked clinically. It is expected that a higher SOP makes prediction more difficult as we go further away from the seizure. Setting the SPH too small may omit important preictal biomarkers whereas making it too large may leak too many interictal patterns which may confuse the model. Besides, different models may provide varying performance with different values of SPH and SOP. Therefore, the practice that we have adapted is to generate benchmarks using multiple combinations using wide ranges of SPH and SOP with the *SPH* ∈ {2, 5, 15, 30} minutes and the *SOP* ∈ {1, 2, 5} minutes. This generates a total of 12 possible combinations of SPH and SOP which will denote our 12 benchmarks as illustrated in Figure 3.

**Fig. 2.**
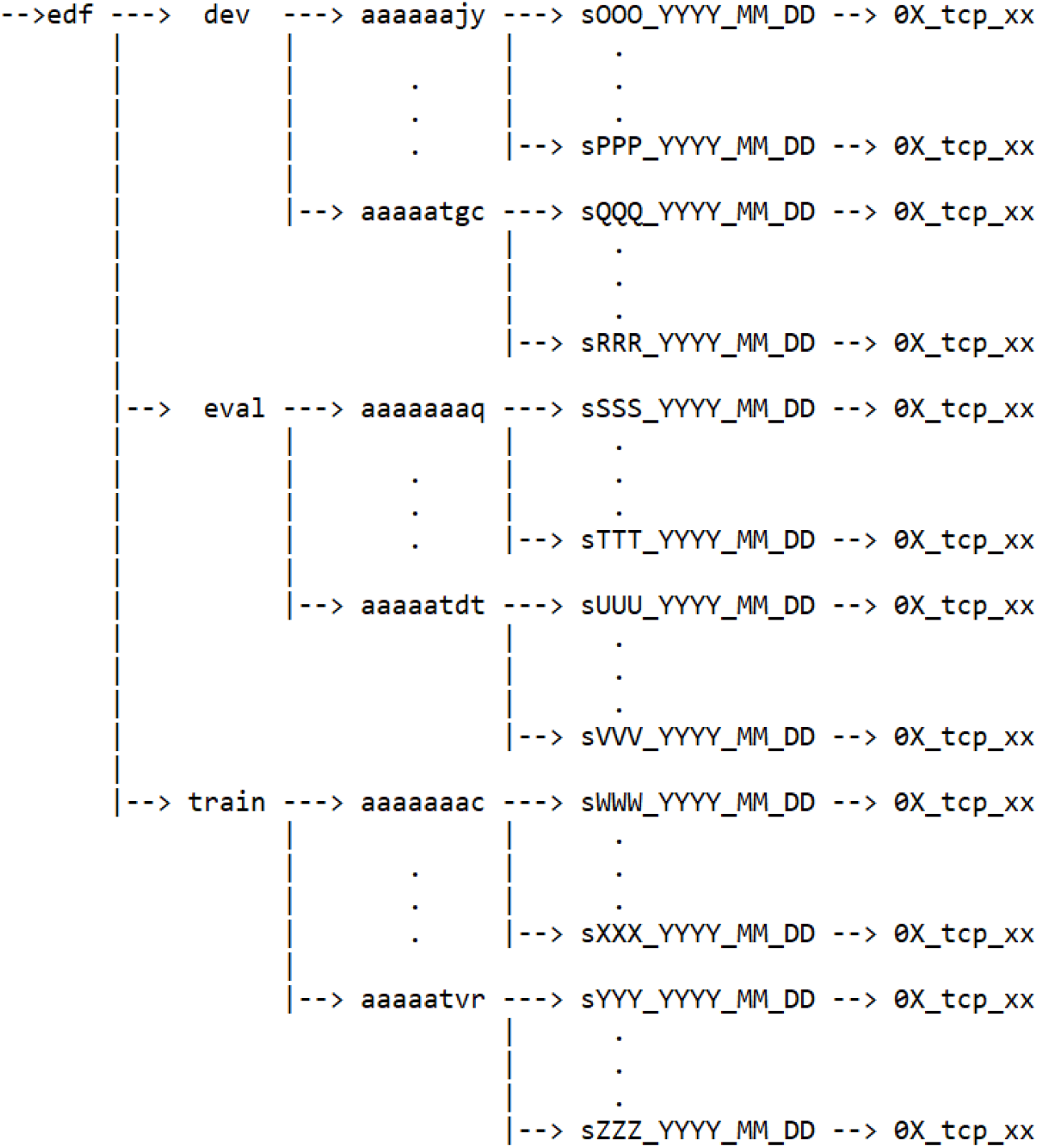
Organization of the TUSZ data.

**Fig. 3.**
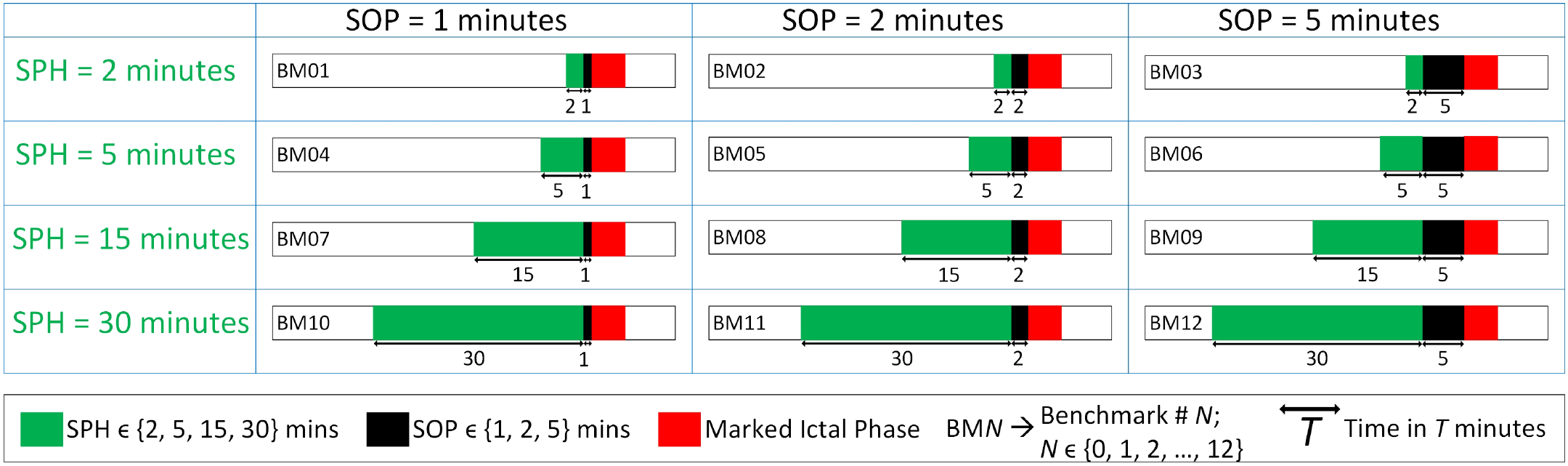
Illustration of the 12 benchmarks in terms of SPH and SOP.

### C. Challenge of Selecting the Correct Window size

Discriminating between preictal and interictal patterns is accomplished using segments of data extracted from long-term EEG. Choosing a segment too small can lead to model underfitting whereas setting it too large may cause it to overfit, increases the data dimensionality quadratically and may not be suitable for time-series classification/prediction. Currently we rely on the state-of-the-art in literature [11], [16] and set *T*_*seg*_ = 5 seconds. However, in future iterations, we plan to identify the benchmarks as *Na, Nb, Nc*, etc… where *N* ∈ {1,, 2,, …, 12} and the characters *a, b, c* represent different values of *T*_*seg*_.

### D. Different Sampling Rates

For our data, the majority of the recordings (5,386/7,377) used a sampling rate of 256 Hz; there were also recordings with 250 Hz, 400 Hz, 512 Hz and 1000 Hz. While such mixed sampling rate data may be a common occurrence in longitudinally collected data, such variation is a critical variable that must be correctly handled. For example, if using raw data as an input with a fixed window size, the number of samples available will be different for each segment along the time-axis. With *T*_*seg*_ = 5, a rate of 250 Hz provides 1250 samples whereas 256 Hz provides 1280 samples per channel. Since ML models are designed to handle a uniform input size, we up/down-sample segments to 256 Hz to make the data uniform with respect to the most commonly occurring sampling rate. Our calculations suggest that the any loss of information as a result of down-sampling is minimal and does not affect the ML model performances.

### E. Different Recording Durations and Seizure Profiles

One of the significant challenges is that each session is of different length and the seizures are spaced out differently. Given that there are more than 4000 recorded ictal events, it requires carefully determining which seizures belong to each benchmark and if enough preictal segments can be extracted. Fig. 4 illustrates this phenomena for each benchmark. Further, it also requires the determination of whether enough interictal segments are available. For some subjects, interictal segments can be extracted from sessions that do not contain seizures but this is not always possible.

**Fig. 4.**
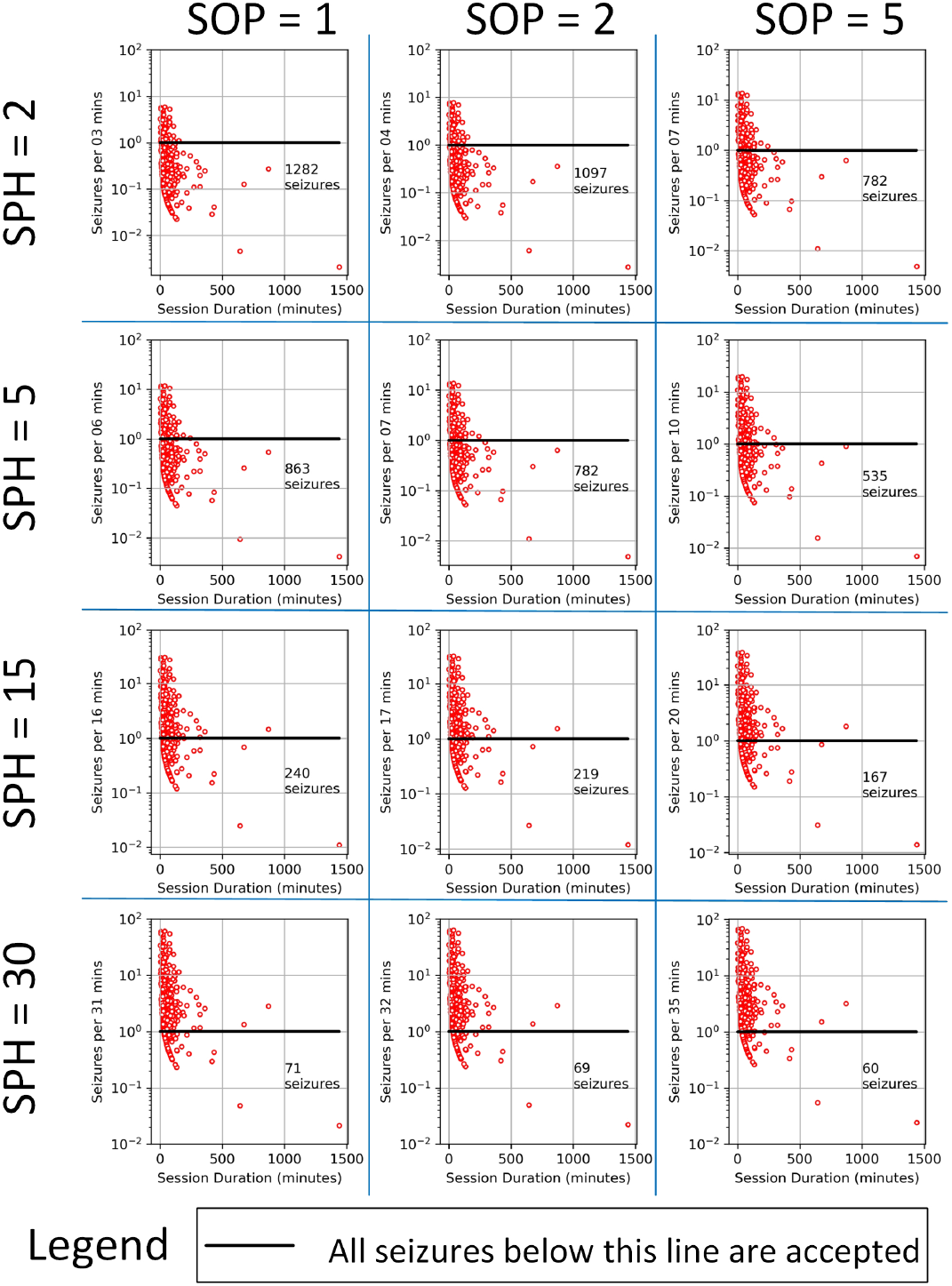
Acceptable seizures for each of the benchmarks. The x-axis shows the distribution of each session whereas the y-axis shows the seizures per unit of gap time *SPH* + *SOP*. Only seizures below the *y* = 1 line are eligible. All times are in minutes.

### F. Other Bottlenecks

These include class imbalance, data scaling, streaming nature and multiple cross-validation folds. Besides the SPH, SOP and *T*_*seg*_, other factors that needs attention are interictal to preictal class imbalance, data normalization, shuffling of validation data and additional pre-processing steps such as filtering. Once all of these design decisions are made, we can build both patient-specific and patient-independent sets which can either be single-fold (a single training, validation and test set each) or multiple-folds for cross-validation (CV) e.g. 5-fold CV. Building N-fold CV datasets allows for the combination of training and validation data. Based on the recommendation of the authors of TUSZ [8], the “dev” (validation) set can be used for tuning model parameters but not the “eval” (test) set which should only be used for final inference/testing. Therefore, we can build N-fold CV sets using data from sessions in both the train and validation directories.

## V. Methods

In this paper, we present 12 benchmarks that are “single-fold” where the interictal segments are randomly sampled, and the data is unfiltered, not normalized and the 5-second segments are non-overlapping. To deal with the above bottlenecks and challenges for creating such ML-ready benchmark datasets, our tool accomplishes the following four tasks:

1. Creation of the metadata
2. Montage Calculation from raw EEG data
3. Seizure profiling and interim dataset creation
4. Creation of the final ML-ready dataset

Hence, in the next few subsections, we will discuss how MLSPred-Bench creates the benchmarks with these four steps.

### A. Metadata Creation

Our major assumption is that in a single session conducted on the same day, EEG data is contiguous and there is no pruning across multiple EDF records. The pruning is only assumed for the same patient across multiple sessions. This is further justified by the date/time information available and our seizure selection process. Therefore, each session is treated as one “subject” from which we will extract preictal and interictal samples associated with each selected seizure and build the benchmarks. In TUSZ, the session metadata is stored in a disjoint manner where there is a unique comma-separated value (CSV) file for each record. To facilitate easier data extraction, we create a single meta-data text file for each session which identifies the number of seizures in each session, and the cumulative start and end times of each seizure. A further advantage is that we designed the format and structure of this text file to match the CHB-MIT summary files, making it easier for users to understand and process.

For fast identification and retrieval, we develop a custom-designed ID convention for each session in the following format: {dtp}_{pid}_s{sid}_{rtp}. The first field comprises three letters that indicate the dataset from which it was taken (training - ‘trn’, testing - ‘tst’ or validation - ‘vld’). The patient ID is identified by the third field. Though the original anonymized ID comprises 8 lower-case English alphabets, only the last three letters are unique; the first 5 are always ‘aaaaa’ for all patients. The third field identifies the session number with a fixed character ‘s’ followed by 3 characters of an integer number in 03d format starting from 001. The last field “rtp” identifies the reference electrode type (either ‘ar1’ - AR1, ‘le2’ - LE2 or ‘ar3’ - AR3). This consolidated session ID is used to name the metadata files, the concatenated raw EEG data and the montage data. For the interim and final ML-ready datasets, the naming convention is described in later sections.

### B. Raw Data and 20-Channel Montages

Creating the concatenated raw data is a straightforward process where data from each record belonging to the same session is added in a continuous order. Using the same convention as earlier, let each EEG record contain *L* samples where

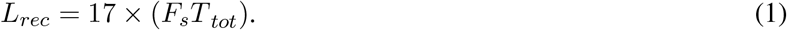

For simplicity, assume *T*_*tot*_ is the same for all records for a particular session and there are *R* records, then the total size of the record will be:

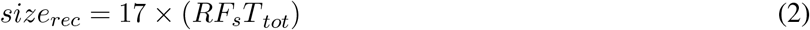

The data from 17 channels is used to calculate the montages as specified in the TUH documents. Let us denote the data from each channel as *{Ch*_0_, *CH*_1_, …, *Ch*_17_}. Further, we are provided the set channel pairs to build the montages as 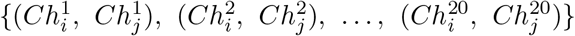 where *i, j* ∈ {1, 2, …, 17} and *i≠ j*. In that case, the content of each montage *k* can be described by the following set of difference equations:

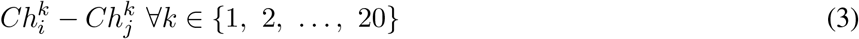

The total size of each montage would then simply be:

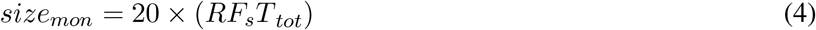

### C. Interim Dataset Extraction

This is the most crucial and complex stage where we use the montages created to extract preictal and interictal data associated with each seizure for each benchmark. This process includes three main stages, filtering out the seizures that belong to each benchmark, extracting the preictal and interictal samples, and up/down-sampling to ensure all sessions have the same input size. Recall that different sampling rates are used across multiple sessions which makes this last step necessary.

#### 1) Isolating Seizures for Each Benchmark

Fig 3 illustrates ictal phase in black and ideally, there will be a pre-ictal phase (green), post-ictal (blue) and large swathes of interictal (red). In the ideal case, the preictal period will align with the SPH which is in the interval [*t*_*start*_ *− SPH − SOP, t*_*start*_ *− SOP*]. Therefore, a seizure can belong to a particular benchmark if and only if no seizure occurs at least *t*_*start*_ *−* (*SPH* + *SOP*) minutes before that seizure. Let us consider two seizures *i* ad *j* with start and end times denoted by *t*_*start,i*_, *t*_*end,i*_ and *t*_*start,j*_, *t*_*end,j*_ where *j* = *i* + 1. Let us also assume a Benchmark *b* with *SPH*_*b*_ and *SOP*_*b*_ ∀*b* ∈ {1, 2, …, 12}. Then, seizure *j* belongs to benchmark *b* if and only if *t*_*start,j*_ *− t*_*end,i*_ ≥ *SPH*_*b*_ + *SOP*_*h*_. We repeat this process iteratively to build up the set of seizures for each benchmark.

#### 2) Extraction of Preictal and Interictal Segments

To facilitate continuous-time seizure prediction, we split the long-term EEG into smaller segments of *T*_*seg*_ seconds and try to identify preictal patterns from these short segments. The total preictal segments for each ictal event will be

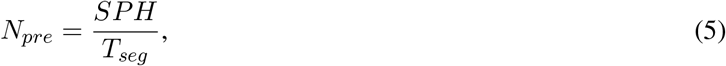

out of a total of:

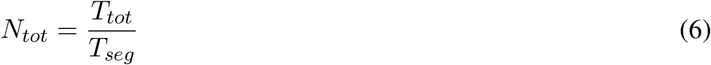

segments. Overall, a segment will contain a total of *L* samples or data points where

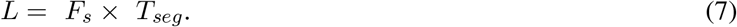

The number of extracted interictal segments *N*_*int*_ is:

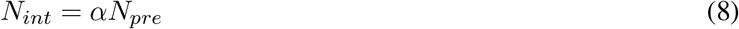

where the ratio of the number of preictal to interictal samples is 1:*α*; *α* ≥ 1 so that we have at least an equal number of interictal and preictal segments to create a balanced dataset (as in this work). Then, there is an equal number of interictal and preictal segments (*N*_*pre*_) for a total of 2*N*_*seg*_ segments associated with each seizure of each benchmark. In general, the number of used segments *N*_*seg*_ is:

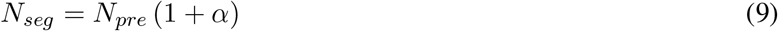

*N*_*seg*_ will vary based on the *SPH* duration as *T*_*seg*_ is currently set to 5 seconds. In general, the data associated with each seizure would be a 3D matrix size 2*N*_*seg*_ *×* 20 *× L*. In other words, there are 2*N*_*seg*_ segments of size 20 *× L* where we have 20 channel montages containing *L* samples along the time-domain. However, because *f*_*s*_ may vary across sessions (though around 80% of the sessions use 256Hz), an extra step is needed to ensure that the *L* is the same for all seizures.

#### 3) Aligning the Number of Samples

The most common sampling rate is 256 Hz and hence, we select it as the base rate. For a 5s window, which will translate to a window size of 5 *×* 256 = 1280 samples. Other than 256 Hz, 4 other sampling rates are observed (250, 512, 400 and 1000 Hz). To keep the pre-processing simple when one of these sampling rates is observed, instead of using re-sampling techniques such as interpolation, we use an approximate method with zero-padding as follows:

- For 250 Hz: we up-sample with zero padding (ZP). Because 5 *×* 250 = 1250, we simply need 30 additional samples to get to 1280. Hence, we add 15 zeros at the start and end of each segment.
- 512 Hz: We simply down-sample to 256Hz by picking 1 out of every 2 samples e.g. odd-numbered.
- 1000 Hz: We down-sample to 250 Hz by picking 1 out of every 4 samples (e.g. 1, 5, 9, …) and then use zero-padding in a similar to the case of 250 Hz.
- 400 Hz: This is the most complicated but also rare case. Here, we down-sample to 250 Hz by picking 5 out of every 8 samples (e.g. {1, 3, 5, 7, 8, …, 1993, 1995, 1997, 1999, 2000} - 5 *×* 400 = 2000 samples) and then use zero-padding similar to 250 Hz.

### D) Creation of the ML-Ready Sets

Once the interim dataset has been created, the data is ML-ready but would still require additional scripting to handle the preictal and interictal samples associated with each seizure. Therefore, we take the additional step of creating a “single-fold” dataset that is ready to be used for ML development. Hence, all preictal and interictal segments associated with seizures belonging to the training set are consolidated into a single “hdf5” file and the labels are consolidated into a single “csv” file for each benchmark. A similar approach is employed for the validation and testing data.

## VI. Data Description and Results

In this section, we provide descriptive results regarding the data itself and ML-based validation results using SPERTL [17]. The naming conventions for the metadata, raw EEG and montages have already been described; we first discuss the final directory structure and naming conventions of the ML-ready data.

### A. Organization and Naming Conventions

Using our method, MLSPred-Bench creates six sub-directories “metadata”, “raweeg”, “montage”, “interim”, “fld_sng” and “fld_mlt_005” in the specified output path to store the ML-ready benchmark datasets. Fig. 5 illustrates the final directory structure. MLSPred-Bench stores the output metadata, concatenated raw EEG data and the calculated montages in the sub-directories ‘metadata’, ‘raweeg’ and ‘montage’, respectively. The interim data which includes the preictal and interictal patterns associated with each seizure for every benchmark are stored in a separate sub-directory called ‘interim’. The single-fold directory called “fld_sng” contains the final dataset for each patient-independent seizure prediction benchmark including training, validation and testing data. In future versions, we intend to make the storage of intermediate data including concatenated raw EEG, montages and interim datasets optional to potentially reduce the storage burden. However, the option to store this data will aid users interested in doing additional pre-processing, feature extraction, patient-specific predictions, etc. and the ability to create ML-ready 5-fold CV sets. Table I describes the final naming convention used for each dataset including example names for the interim and single-fold datasets.

**TABLE I.**
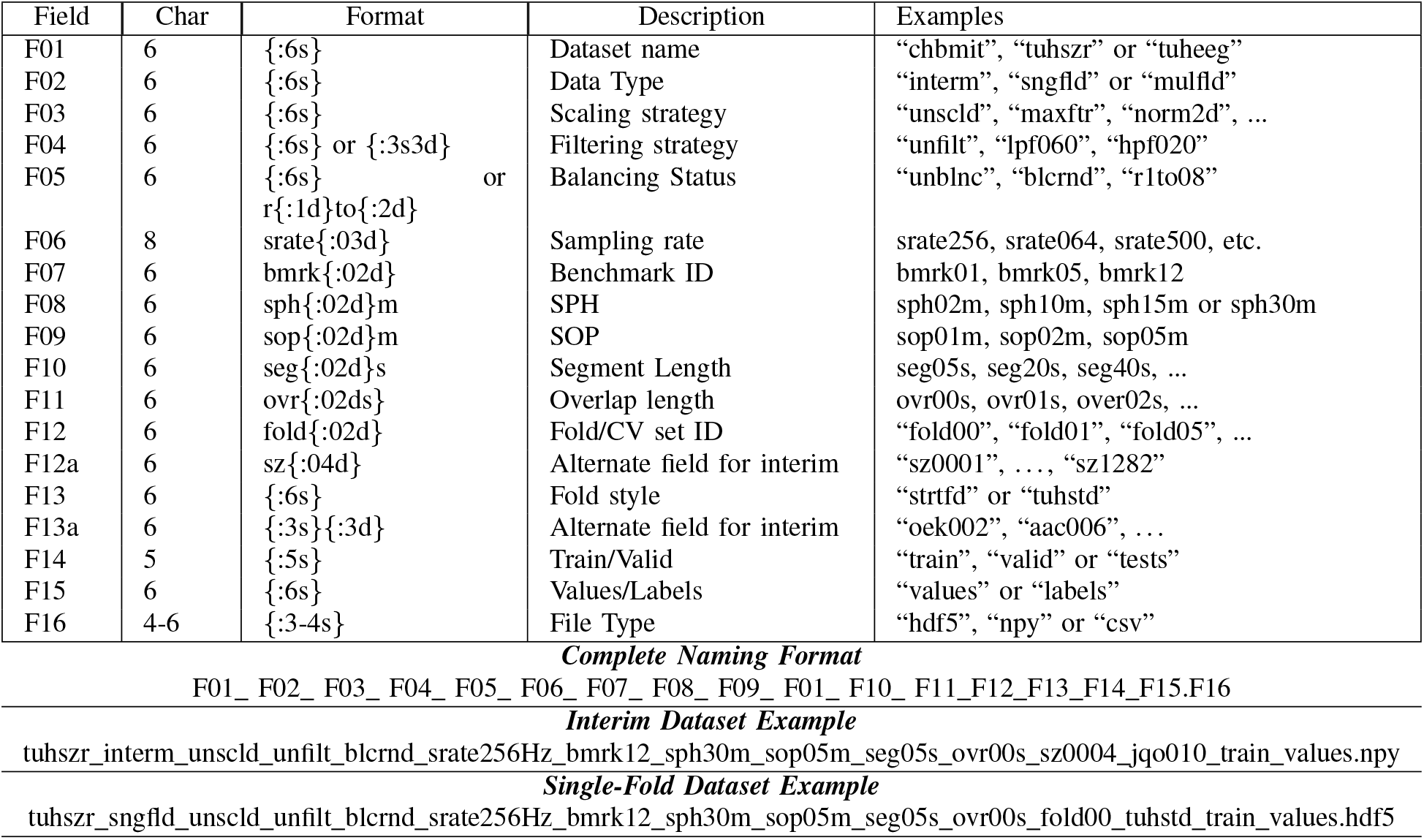
Naming Conventions.

**Fig. 5.**
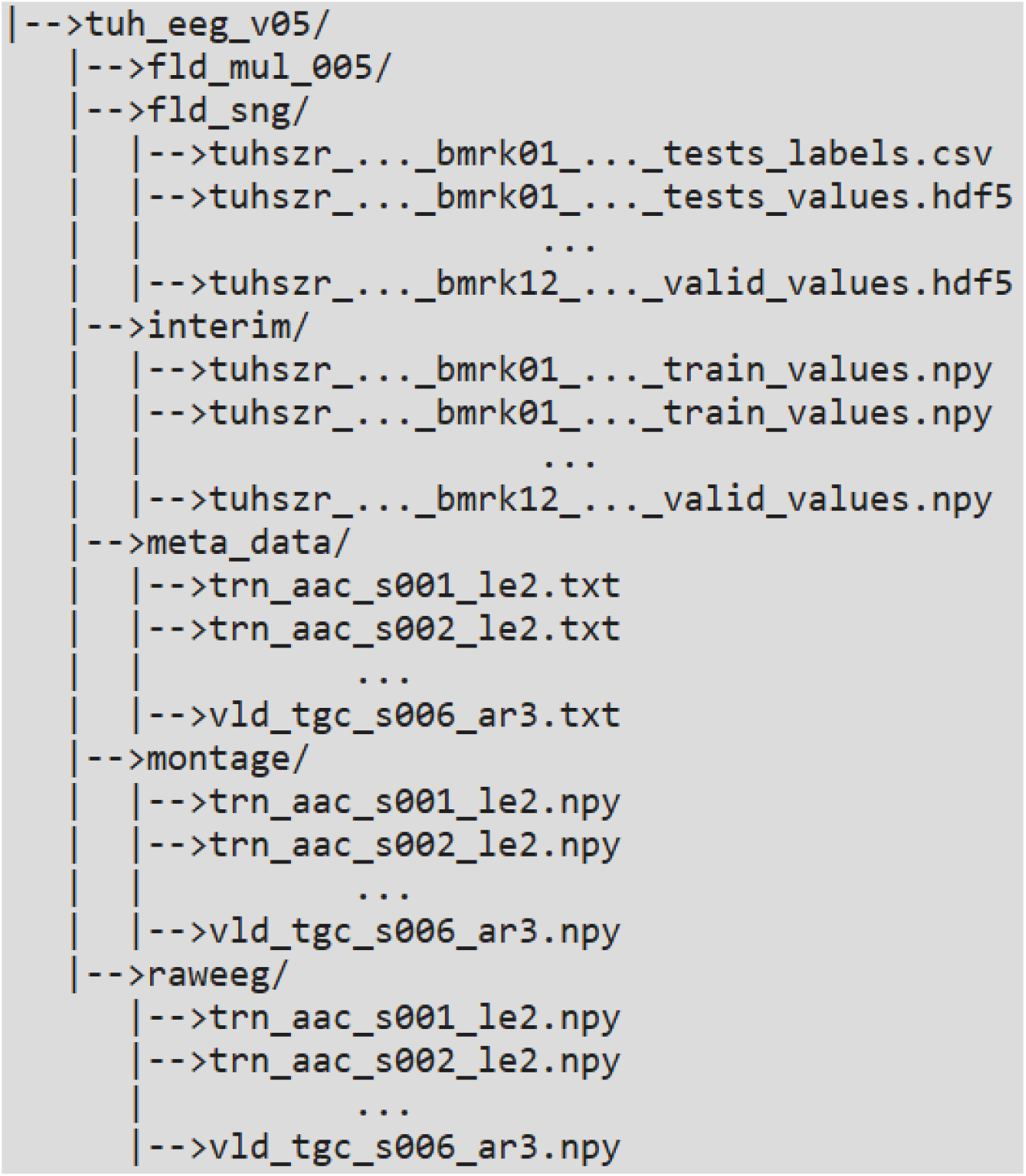
Directory Structure of the final data generated. There are separate directories each for raw data, montages and the interim dataset. The ML-ready data is in the ‘fld_sng’ directory. An empty directory is left for future use with 5-fold cross-validation sets.

Fig. 6 illustrates the number of seizures that belong to each benchmark. We note that as we increase the required gap time defined by *SPH* + *SOP*, fewer seizures are available for benchmarking. We suspect that one of the major reasons for this is the pruning of long-duration segments with no seizures. However, another factor is the observance of seizure clusters in certain patients and this phenomena is observed in multiple datasets. The train-valid-test distribution follows the seizure distributions in the interim dataset and are provided in Table II. It can be observed that the size of the training set roughly ranges between 50-60% whereas the validation set is as large as 33%. There is a chance that this may result in an inadequate amount of training data.

**TABLE II.** Train-Valid-Test Split.

| Benchmark (gap) | Train | Valid | Test |
| --- | --- | --- | --- |
| BM01 (3 mins) | 53.59% | 32.84% | 13.57% |
| BM02 (4 mins) | 52.14% | 33.73% | 14.13% |
| BM03 (7 mins) | 52.26% | 33.02% | 14.72% |
| BM04 (6 mins) | 52.81% | 32.23% | 14.96% |
| BM05 (7 mins) | 52.15% | 30.65% | 17.20% |
| BM06 (10 mins) | 52.92% | 30.00% | 17.08% |
| BM07 (16 mins) | 53.88% | 30.59% | 15.53% |
| BM08 (17 mins) | 55.09% | 28.74% | 16.17% |
| BM09 (20 mins) | 58.97% | 26.92% | 14.10% |
| BM10 (31 mins) | 59.15% | 26.76% | 14.08% |
| BM11 (32 mins) | 57.97% | 27.54% | 14.49% |
| BM12 (35 mins) | 60.00% | 26.67% | 13.33% |

**Fig. 6.**
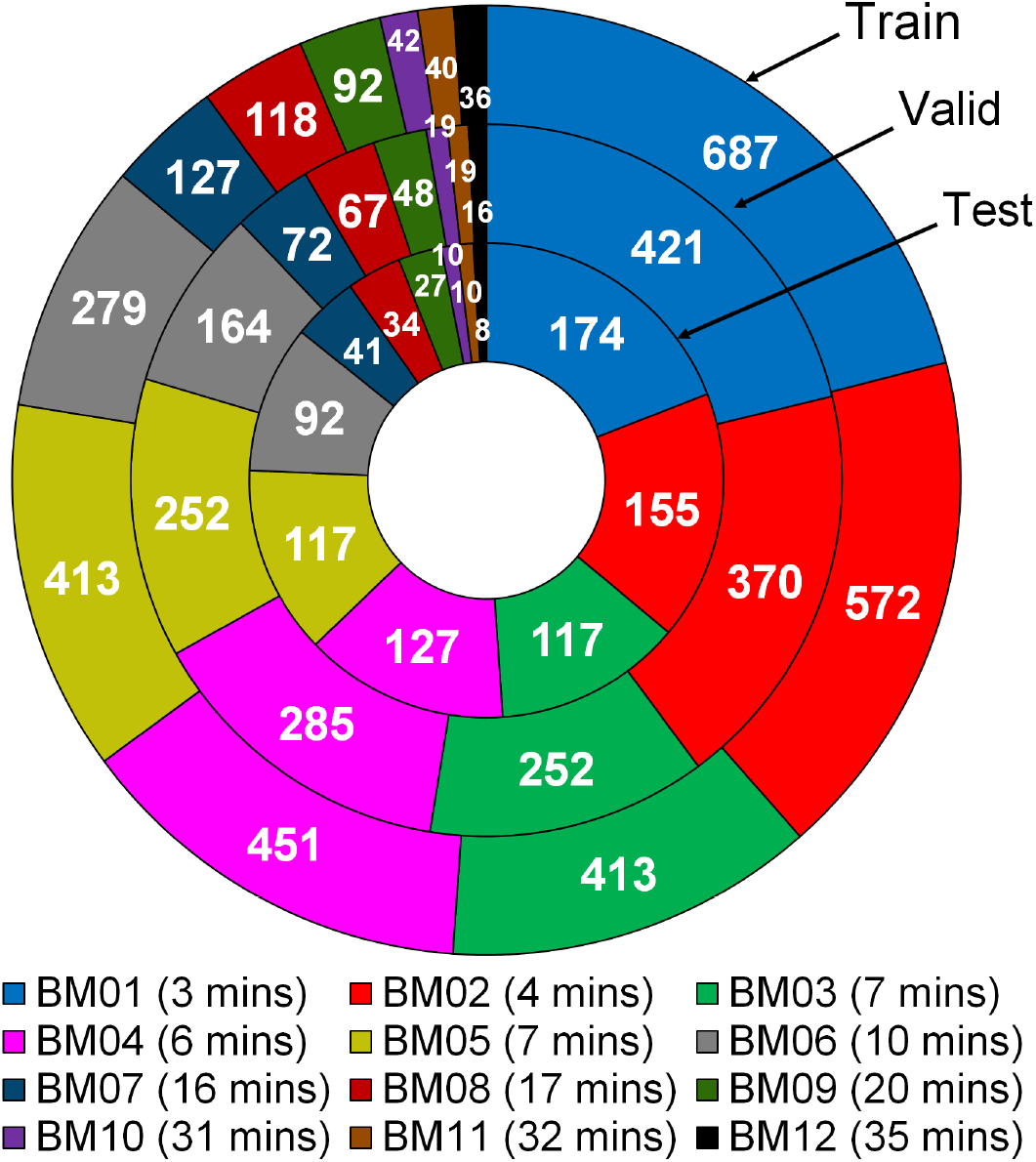
Seizure distribution among the different benchmarks.

### B. Technical Validation

In this section, we modified our previously published SPERTL architecture [17] and trained and tested on all benchmarks. The architecture is based on the residual neural networks (ResNets) and features an input convolutional block followed by 4 residual blocks. Each convolutional block contains a 7×7 convolution filter, batch normalization, dropout (75%), 2×2 max pooling ReLU activation. Each residual block contains 2 convolutional blocks with a skip connection from the previous convolutional block right after the convolution filter. The last residual block is followed by 2 dense layers and a soft-max layer for classification. We used a learning rate of 1e-3, 10% reduction on plateau and early stopping with a maximum of 25 epochs.

Fig 7 illustrates the obtained receiver-operating-characteristic (ROC) curves for each benchmark. In general, it is observed that the model is able to provide a high area under the curve (AUC) score for training which means the dataset is trainable. For some benchmarks (7, 8, 9 and 11), the ROC-AUC score is in excess of 0.79 with benchmark 11 providing the highest score of 0.925. This indicates that the dataset is trainable and with more hyperparameter tuning for each benchmark, we should be able to considerably improve the ROC-AUC score. Although the validation performance is lower, it is comparable to the seizure detection accuracy achieved by the TUSZ authors [8]. Therefore, this validates our methods for creating the benchmarks and implies our tool would be indispensable for researchers interested in seizure prediction.

**Fig. 7.**
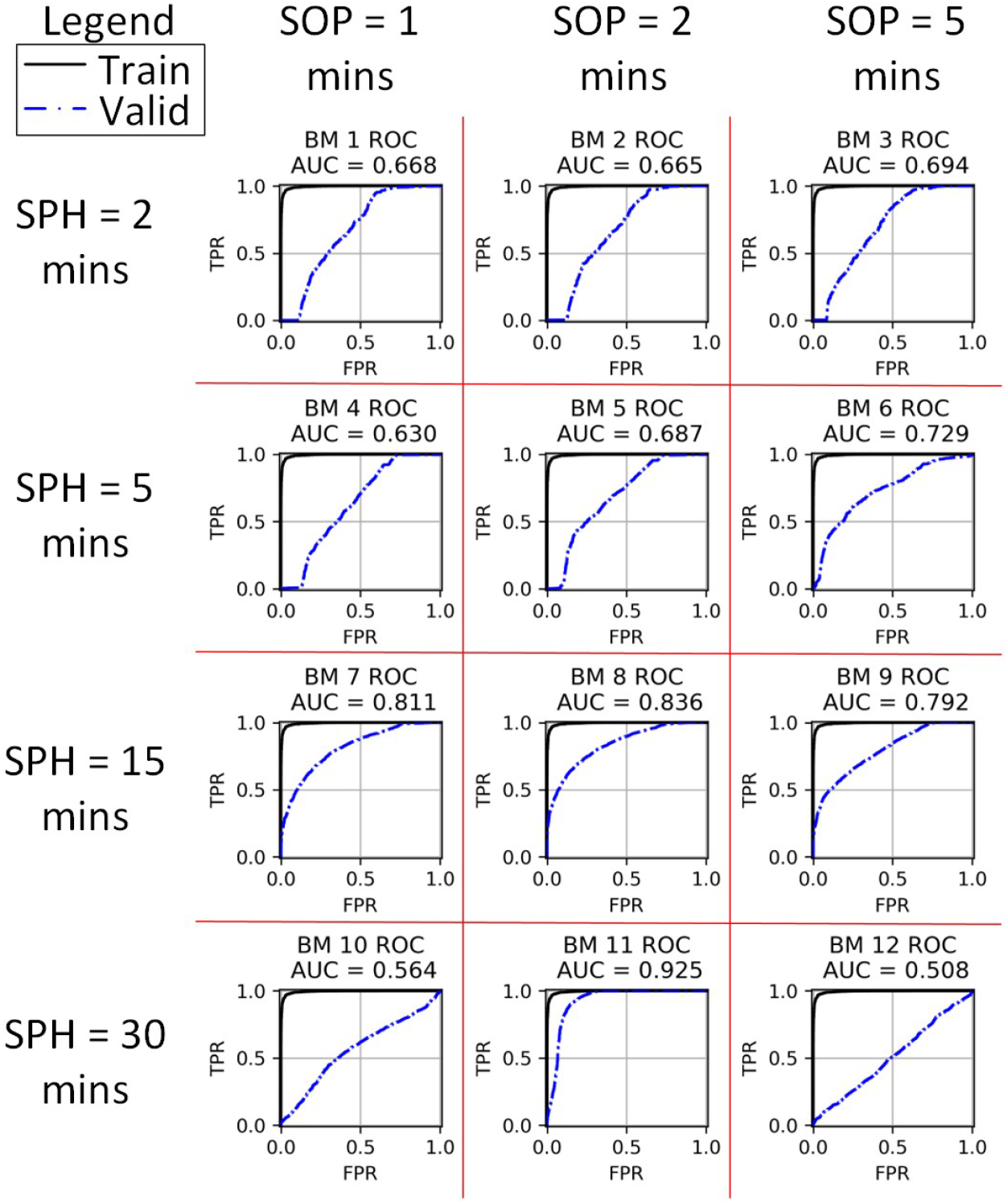
Training and validation results from SPERTL after re-training for the TUSZ patient-independent seizure prediction task.

### C. Code Availability and Usage

The code is available in the following GitHub repository: https://github.com/pcdslab/MLSPred-Bench. The authors assume that users are able to independently obtain the latest version of the TUSZ data and install the packages specified by the “requirements.txt” file.

## VII. Discussions and Conclusion

In this paper, we designed and developed a strategy that results in ML-ready data suitable for modelling epileptic seizure prediction. To demonstrate our strategy, we re-purposed an existing EEG dataset that has been annotated for seizure detection - into an ML-ready seizure-prediction data set. Using the TUSZ dataset we demonstrated the generation of 12 ML-ready seizure prediction benchmarks based on a combination of SPH and SOP duration. As a result of this effort, more than 500G of data was generated which could be used by developers for predictive models. To demonstrate that the resulting dataset is suitable for developing predictive models by showcasing it using the existing model SPERTL. We believe that the resulting benchmarking dataset will enable a single point of reference for machine-learning practitioners, providing a fair way to compare predictive models.

We intend to improve on these benchmarks by taking the following steps: The 5-second windows need to be supplemented with other sizes with an option for overlap. These would allow for time-aligned scoring such as those in [5]. One limitation of following TUSZ convention is an uneven distribution of the training and validation data across benchmarks. Therefore, smarter strategies which can also be used to create multiple cross-validation folds (as is the standard for ML model development) need to be explored. Several other improvements can be made such as testing for different ratios of class-imbalance, assuming a streaming flow of data whereas random distribution is assumed currently, and more pre-processing steps such as filtering can be incorporated.

We strongly believe that the resulting tool will have wide variety of usage of re-purposing detection-annotated data into prediction-annotated data. The repurposed data sets will enable rapid ML-prototyping, and provides benchmarking datasets suitable for fair comparison across different ML models. We encourage the scientific community to use it for their ML models, and contribute in the open-source code to improve the resulting benchmarks.

---

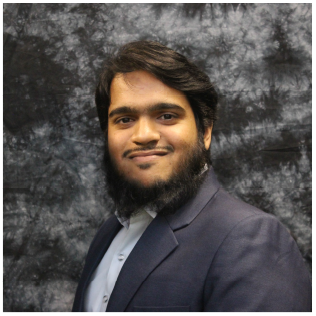

**Umair Mohammad** (S’12–M’20) is a Postdoctoral Associate at Florida International University (FIU) in the Knight Foundation School of Computing and Information Sciences (KFSCIS). Umair earned his PhD in Electrical Engineering from the University of Idaho in 2020. His current research interests include edge machine learning (ML), mobile edge computing, and ML for biomedical event prediction. Umair is particularly interested in early epileptic seizure prediction, developing suitable datasets and advanced deep learning for improving prediction sensitivity.

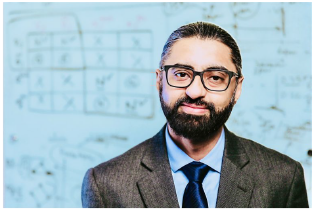

**Fahad Saeed** is a Tenured Associate Professor at KFSCIS and Associate Professor (Courtesy) of Human and Molecular Genetics at the Herbert Wertheim School of Medicine. He earned his PhD from the University of Illinois at Chicago in 2010. Dr. Saeed is a Senior Member of the IEEE and ACM, and he is a National Science Foundation (NSF) CRII and CAREER Award Winner. He has over 90 published peer-reviewed articles, 2 book chapters, multiple patents, edited 3 special issue journals, 4 conference proceedings and 1 book. His current research interests lie at the intersection of high-performance computing and machine learning for proteomics, image-based diagnostics and biomedical event detection.

